# Tumor spheroid elasticity estimation using mechano-microscopy combined with a conditional generative adversarial network

**DOI:** 10.1101/2024.04.12.589174

**Authors:** Ken Y. Foo, Bryan Shaddy, Javier Murgoitio-Esandi, Matt S. Hepburn, Jiayue Li, Alireza Mowla, Danielle Vahala, Sebastian E. Amos, Yu Suk Choi, Assad A. Oberai, Brendan F. Kennedy

## Abstract

Techniques for imaging the mechanical properties of cells are needed to study how cell mechanics influence cell function and disease progression. Mechano-microscopy (a high-resolution variant of compression optical coherence elastography) generates elasticity images of a sample undergoing compression from the phase difference between optical coherence microscopy (OCM) B-scans. However, the existing mechano-microscopy signal processing chain (referred to as the algebraic method) assumes the sample stress is uniaxial and axially uniform, such that violation of these assumptions reduces the accuracy and precision of elasticity images. Furthermore, it does not account for prior information regarding the sample geometry or mechanical property distribution. In this study, we investigate the feasibility of training a conditional generative adversarial network (cGAN) to generate elasticity images from phase difference images of samples containing a cell spheroid embedded in a hydrogel. To train and test the cGAN, we constructed 30,000 elasticity and phase difference image pairs, where elasticity images were generated using a parametric model to simulate artificial samples, and phase difference images were computed using finite element analysis to simulate compression applied to the artificial samples. By applying both the cGAN and algebraic methods to simulated phase difference images, our results indicate the cGAN elasticity images exhibit better spatial resolution and sensitivity. We also evaluated the cGAN on experimental phase difference images of real spheroids embedded in hydrogels and compared the cGAN elasticity with the algebraic elasticity, OCM, and confocal fluorescence microscopy, and found the cGAN elasticity is often more robust to noise, especially within stiff nuclei.

## 1 Introduction

Biological cells sense mechanical stimuli (i.e., applied forces) and convert these to biological responses via a process known as mechanotransduction [1], whereby the mechanical properties of both the extracellular matrix and the cells themselves impact cellular proliferation [2], migration [3,4], and differentiation [2,4,5], as well as the progression of diseases, such as cancer [6] and cardiovascular disease [7]. While early studies predominantly measured the mechanics of two-dimensional (2D) cell cultures, more recently, the focus has shifted to three-dimensional (3D) cultures, which provide more physiologically-relevant data [8]. For example, spheroids (3D clusters of cells that provide both cell-cell and cell-matrix interactions [9,10]) can be cultured in hydrogels, which results in an extracellular matrix-like environment with customizable mechanical and biochemical properties [11].

Initially, techniques for imaging nano-to micro-scale mechanical forces and properties were restricted to two lateral dimensions, however, the rise of 3D cell cultures has spurred the development of 3D imaging capabilities [12]. One technique is traction force microscopy, in which the tractions exerted by cells on their surrounding environment is measured in 2D or 3D [13–15]. Techniques that can measure mechanical properties include atomic force microscopy [16], Brillouin microscopy [17,18] and optical coherence elastography [19,20]. Atomic force microscopy can provide sub-nanometer lateral resolution images of Young’s modulus [21], however, it is usually limited to 2D imaging of the sample surface [16]. Brillouin microscopy can provide sub-micron resolution of longitudinal modulus [22,23], however, the Brillouin signal depends strongly on water content [24,25] and has a penetration depth of ∼100 μm in highly scattering media, which may be insufficient for imaging entire spheroids [26]; furthermore, long acquisition times (up to ∼1 s per pixel) and high-powered lasers are often required, which may damage the sample [17].

In compression optical coherence elastography (OCE), the sample is compressed and the axial displacement is measured, usually from the phase difference between OCT scans [19]. Calculating the gradient of displacement over depth (e.g., by linear fitting) yields the axial strain throughout the sample [27,28]. In a variant of compression OCE called quantitative micro-elastography [29], by compressing a compliant silicone layer with a known stress-strain curve against the sample, the axial stress at the layer-sample interface is calculated from the strain in the layer [30] and Young’s modulus is determined by dividing the axial stress by the axial strain [29]. Using this technique, Hepburn et al. demonstrated the capability to map the elasticity of adipose-derived stem cells embedded in hydrogels [31], while Mowla et. al. developed a variant, mechano-microscopy, that uses optical coherence microscopy (OCM) to obtain sub-cellular elasticity resolution (5 μm) [11,32,33]. However, this technique assumes a frictionless mechanical model where the stress is uniaxial (i.e., acting only along the depth axis) and axially uniform (i.e., the stress at some location within the sample equals the stress at the layer-sample interface directly above or below that location). In practice, these assumptions are often violated by shear stresses caused by conditions at the sample boundary (e.g., friction [34] and surface roughness [35,36]) as well as mechanical heterogeneity within the sample, which reduces the accuracy and precision of elasticity measurements [34,35,37]. Furthermore, in signal processing, errors from phase unwrapping algorithms to convert phase difference to displacement cause artefacts in elasticity images; also, spatial resolution is limited by the fitting range used to compute strain from displacement [28,38,39].

The central problem is how to determine the sample mechanical properties, given the sample displacement and boundary conditions. This is known as the inverse problem [20,40,41], for which solutions have been presented in ultrasound elastography [42]. In OCE, Dong et. al. implemented iterative solutions to the inverse problem in 2D [40] and 3D [20] in which the shear modulus (which is proportional to Young’s modulus for incompressible materials) throughout the sample is initially guessed, the forward problem is solved using finite element analysis (FEA) to obtain the simulated displacement, and the error between the measured and simulated displacement is minimized by iteratively updating the shear modulus and re-solving the forward problem. A main advantage is, unlike in quantitative micro-elastography, the assumptions of uniaxial and axially uniform stress in the sample are not required, thus potentially enabling more accurate and precise elasticity measurements; additionally, spatial resolution may be improved since fitting to compute strain is not required. However, this approach is computationally demanding, requiring ∼20 hours on a high-end desktop computer to solve a typical 3D problem [20]. Furthermore, it does not provide a convenient mechanism to embed prior knowledge about the spatial distribution of the mechanical properties of the sample, and since the algorithm is deterministic, it does not provide an estimate of the uncertainty in its prediction.

One promising method to avoid these challenges is via generative adversarial networks (GANs), which are a machine learning framework that provide a means of learning deep representations of data [43]. GANs consist of a pair of networks (often called the generator, and the discriminator or critic) trained in competition with each other [43]. Popular uses for GANs in medical imaging include artefact removal, cross-modality medical image synthesis, and segmentation [44]. While the input to a GAN is typically a random vector (termed the latent vector), more recently developed conditional GANs (cGANs) also accept a single instance of another random vector (which can encode prior knowledge) and generate images that are conditioned on this vector [45]. For example, in ultrasound elastography, cGANs have been used to synthesize elasticity given prior knowledge of the sample structure and/or displacement [46–48]. The problem of spheroid elasticity estimation with micro-scale resolution is particularly suited to cGANs since, at this scale, spheroids can be represented parametrically (e.g., in terms of the number of cells, cell size, and nucleus elasticity), thus enabling the training set to be generated by simulation using FEA. This enables the use of mechanical models that do not assume uniaxial or axially uniform stress, allows prior knowledge of the spheroid structure to be incorporated, and avoids the significant hurdle of assembling the training set from experimental data. Other advantages presented by cGANs for spheroid elasticity measurement include offline training (avoiding lengthy computations during elasticity generation); direct mapping of phase difference to elasticity, thus avoiding artifacts from phase unwrapping and resolution reduction caused by the strain fitting range; and uncertainty quantification by computing statistics using an ensemble of output images that are generated for a single measurement vector. However, despite these advantages, cGANs have not yet been applied to spheroid elasticity estimation, or indeed to OCE more generally.

In this study, we investigate the feasibility of using cGANs to synthesize elasticity maps of tumor spheroid samples conditioned upon phase difference. Using a parametric spheroid model, we generated 5,000 artificial spheroid elasticity maps, which were used as input to an FEA solver to generate simulated elasticity/displacement image pairs. To avoid the need for phase unwrapping, the simulated displacement images were converted to phase difference images. Noise was added to the phase difference images using a model that incorporates the dependence of phase difference noise on the OCT SNR, which varies with depth due to the OCT confocal function. The cGAN was trained using pairwise samples of simulated elasticity images and noisy phase difference images. To evaluate the cGAN, we compared the elasticity generated by the cGAN using the simulated phase difference to the known ground truth elasticity, as well as to elasticity generated using the existing processing chain for mechano-microscopy, referred to as the algebraic method (see Sec. 2.1.1). Additionally, we measured the phase difference in real tumor spheroid samples using mechano-microscopy and compared the elasticity generated by the cGAN using these measurements to that computed using the algebraic method, as well as with co-registered OCM and confocal fluorescence microscopy (CFM) images. Our results show that cGANs can generate improved elasticity images compared to the algebraic method while additionally enabling quantification of the prediction uncertainty based on the ensemble of elasticity images generated for each phase difference image. In very stiff regions (e.g., within nuclei), the algebraic method often provides negative elasticity due to a breakdown of the mechanical model and phase noise, which is not physically meaningful, while the cGAN often produces high elasticity. Overall, our results indicate that the cGAN elasticity images are more robust to noise and deviations from a uniaxial compressive model than the algebraic elasticity images.

## 2 Method

### 2.1 Elasticity image processing

Here, we provide a brief overview of the algebraic method [29,32,33], followed by a description of the methods and potential benefits associated with using cGANs.

#### 2..1.1 Algebraic method

In mechano-microscopy, the elasticity throughout a sample is computed by compressing the sample and acquiring OCM B-scans (2D cross-sectional images) both before and after compression. The Young’s modulus, 𝐸, is

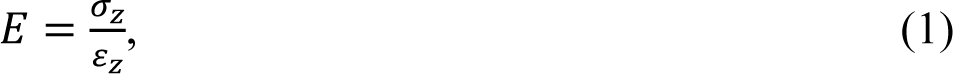

where 𝜎_𝑧_ is axial stress and 𝜀_𝑧_ is axial strain measured in a homogeneous material undergoing uniaxial compression or tension [29]. In mechano-microscopy, the 𝑧 axis is parallel to both the OCM optical axis and the compression axis. The axial stress, 𝜎_𝑧_, is determined from the deformation of a compliant silicone layer that is compressed with the sample, for which the stress-strain curve has been pre-characterized [29,30]. While this method allows for variation in 𝜎_𝑧_ along the B-scan lateral axis (denoted the 𝑥 axis) since the layer deformation can vary along the 𝑥 axis, it assumes 𝜎_𝑧_ is uniform along the 𝑧 axis; furthermore, errors in the measurement of 𝜎_𝑧_ can result from many factors including air gaps between the layer and the sample, friction at the layer boundaries, and inaccurate detection of the layer-sample interface [34,35,39,49].

Axial strain, 𝜀_𝑧_, is calculated by measuring the axial displacement, 𝑢_𝑧_, due to compression at every (𝑥, 𝑧) location and computing the gradient of 𝑢_𝑧_ with respect to 𝑧, i.e., 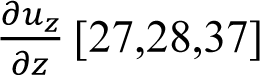. Methods for computing the gradient include linear regression [27,28] and finite difference [50] methods; however, the presence of noise typically necessitates increasing the fitting range or performing spatial averaging to improve sensitivity, at the expense of resolution [38,39].

Axial displacement is measured from the phase difference, Δ𝜙, between the complex OCM B-scans before and after compression using [27,28]

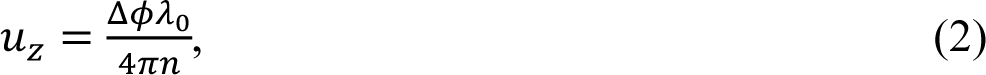

where 𝑛 = 1.4 is the assumed refractive index of tissue [51], and 𝜆_0_is the central wavelength of the light source. Using the top of the image as the reference point (i.e., where zero displacement occurs), for a compressed sample, we expect displacement to generally decrease with 𝑧, where negative displacement corresponds to movement towards the reference. However, since phase difference is limited to the range (−𝜋, 𝜋], phase difference images often exhibit phase wrapping, where the phase difference returns to 𝜋 instead of decreasing past −𝜋 . Previously, phase unwrapping algorithms have been used to convert phase difference to displacement [27,28,32], however, these algorithms often produce artifacts, particularly in noisy regions of the image [28].

Finally, elasticity images are generated by dividing the axial stress by the axial strain at every location in the B-scan. While this is a fast and simple method of estimating elasticity, it assumes stress is uniaxial and axially uniform; as such, the presence of significant shear stresses may produce artifacts. For example, this method will yield a negative Young’s modulus (which is unphysical) if the calculated stress and strain have opposite signs, which may occur due to shear stresses. Therefore, such regions are often masked out of the final elasticity image [52].

#### 2.1.2 cGAN method features

As an alternative to the algebraic method, we propose to train a cGAN to learn the correspondence between OCM phase difference images and elasticity images; that is, given the input of a phase difference image created using mechano-microscopy and a latent vector, the cGAN should output an elasticity image instance from the learned distribution conditioned upon the phase difference. This method presents several advantages compared to the algebraic method [29,32,33] and iterative methods [20,40], as outlined below.

##### 2.1.2.1 Inclusion of complex physics and prior knowledge

A key advantage of the cGAN method is that it enables the incorporation of complex physics and prior knowledge of the sample structure by using simulated data from FEA to train the cGAN. Importantly, FEA can account for the presence of shear stresses and strains, which is a source of negative elasticity in the algebraic method, as well as variations in axial stress with depth, thus enabling a more accurate representation of the sample. Furthermore, prior knowledge of the sample structure (e.g., the fact the sample contains one spheroid, consisting of tens of cells, each containing one relatively stiff nucleus, all of which is embedded in hydrogel) can be included in the training set by generating artificial samples using a parametric model that incorporates this information. Generating the training set by simulation also enables a much larger and more diverse training set to be constructed, compared to using real experimental data, which is critical for training the cGAN. The cGAN method also does not require the compliant layer that is used to determine the axial stress in the algebraic method, thus avoiding the errors associated with air gaps, friction, and erroneous detection of the layer-sample interface. However, as the inverse elasticity problem is ill-posed, (i.e., a given phase difference image could correspond to a range of valid elasticity images), it should be noted that to overcome ambiguity, the elasticity of at least one component in the image (e.g., the hydrogel) must be known.

##### 2.1.2.2 Processing speed

While the algebraic method enables rapid elasticity image generation, this comes at the cost of the accuracy of the solution, since the rapid processing is made possible by making simplifying assumptions such as uniaxial and axially uniform stress. Conversely, while iterative methods enable improved solution accuracy, long compute times are required. For the cGAN method, while a large upfront computational cost is required to perform the FEA simulations and train the cGAN, once trained, cGANs enable rapid elasticity image generation while incorporating complex physics and knowledge of the sample structure to improve accuracy.

##### 2.1.2.3 Uncertainty quantification

Since the cGAN learns a distribution of elasticity images conditioned upon the phase difference images, the precision of the cGAN elasticity image for each measurement can be estimated by varying the latent vector to generate an ensemble of elasticity images, then calculating the standard deviation, which aids the interpretation of cGAN elasticity images by providing important information on the reliability of the cGAN elasticity estimate at each spatial location.

As the algebraic and iterative methods are deterministic, standard deviation images cannot be generated. However, the precision of the elasticity measurements may be estimated by characterizing the elasticity sensitivity based on the system noise and signal processing chain [39], although this usually does not provide spatially-resolved maps of precision.

##### 2.1.2.4 Direct analysis of phase difference

Another advantage of cGANs is that they can be trained to learn the direct correspondence between phase difference images and elasticity images. This avoids needing to perform phase unwrapping to produce displacement images in the algebraic and iterative methods, which can produce artefacts [28]. It also avoids the need for strain estimation, which is necessary in the algebraic method. Therefore, spatial resolution may be improved using the cGAN method, since linear regression or spatial averaging for strain estimation is not needed.

### 2.2 Architecture of the cGAN

The cGAN architecture was based on a structure previously described by Ray et al [47]. Briefly, the cGAN consists of two parts, a generator, and a critic. The generator learns to produce a distribution of elasticity images corresponding to an input phase difference image, while the critic pushes the generator to produce better images. The generator inputs are a phase difference image and a 100-dimensional latent vector, and the generator output is a corresponding elasticity image instance from the learned conditional distribution, where additional instances are generated by varying the latent vector. All images are 256 × 256 μm^2^ (256 × 256 pixels) corresponding to the OCM B-scan plane. The generator is based on the popular U-Net architecture [53], which excels at image-to-image transformations. As shown in Figure 1a, the generator consists of a contracting path (in which the image is progressively downsampled to enable feature extraction at different scales) followed by an expanding path (in which the image is progressively upscaled to achieve the desired output dimensions), with skip connections that transfer learned features from the contracting path to the expanding path. Conditional instance normalization is used to inject the latent vector at each scale of the generator, as outlined by Ray et al [47].

**Figure 1.**
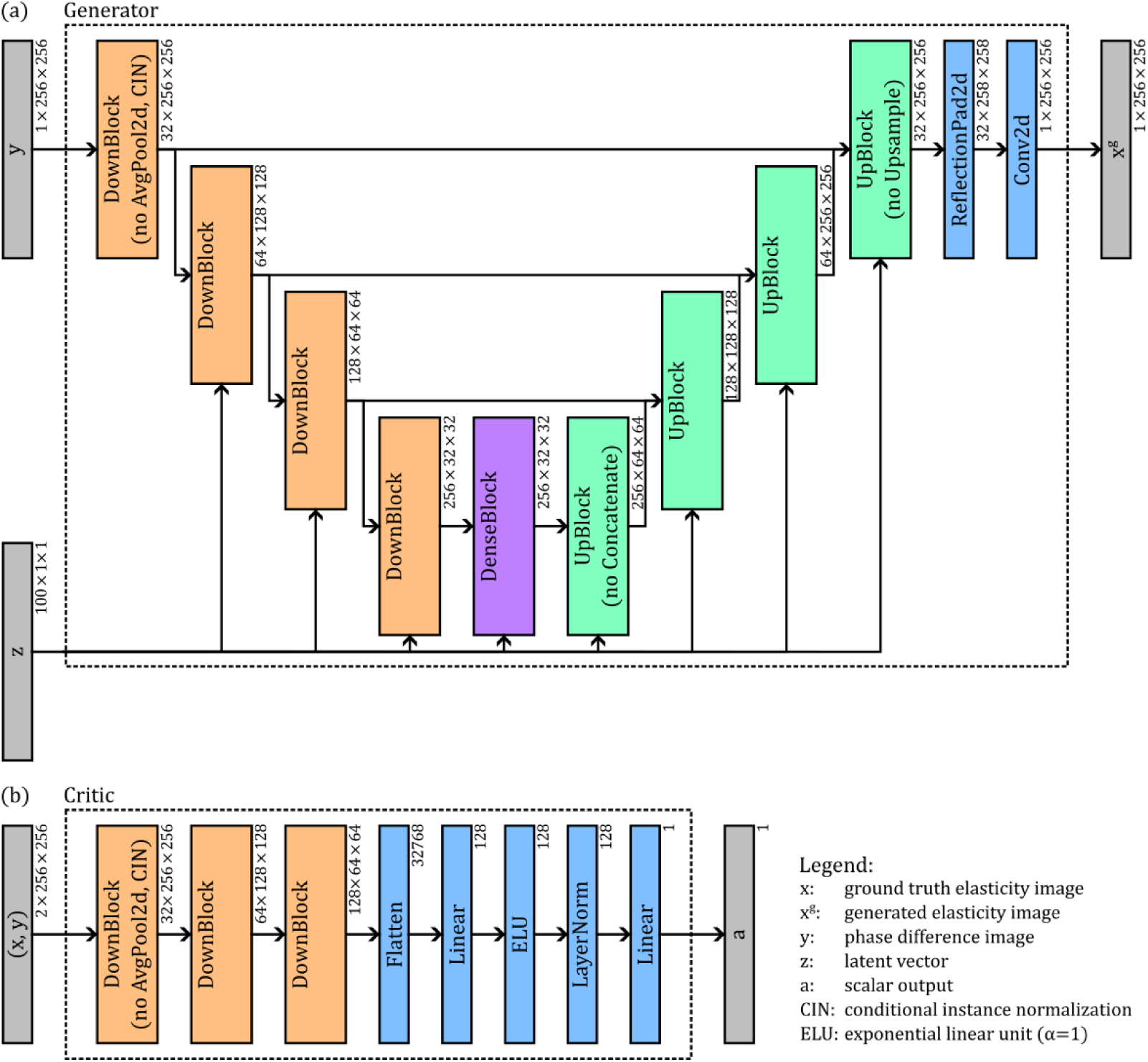
Architecture of the cGAN showing the arrangement of the main building blocks (i.e., the DownBlock, DenseBlock, and UpBlock) of the generator and critic networks.

The critic inputs are the paired phase difference and elasticity images, and the output is a scalar indicating whether the elasticity image is from the generator. As shown in Figure 1b, the critic uses a simple convolutional neural network architecture consisting of two downsampling layers followed by two fully connected layers.

The loss functions for the generator and critic are defined such that the critic is trained to differentiate ground truth elasticity images (produced artificially, as described in Sec. 2.3) from the elasticity images produced by the generator, while the generator is trained to fool the critic by producing elasticity images that resemble the ground truth elasticity images. The loss function, ℒ(𝑑, 𝑔), is described in detail by Ray et al [47] and shown in Eq. 3,

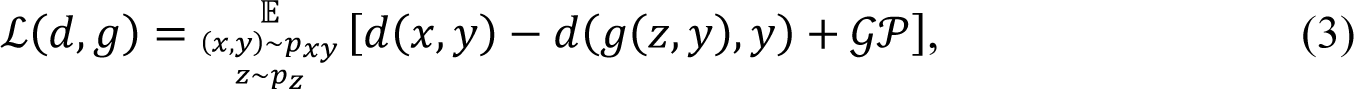

where 𝑑(𝑥, 𝑦) is the output from the critic; 𝑔(𝑧, 𝑦) is the output from the generator; (𝑥, 𝑦) represents an elasticity/phase difference image pair from the true distribution, 𝑝_𝑥𝑦_; 𝑧 represents the latent vector from the latent space, 𝑝_𝑧_ ; 𝒢𝒫 is the gradient penalty term used to enforce Lipschitz continuity on the critic, and 𝔼 denotes the expectation value. Training the cGAN thus involves solving the min-max problem,

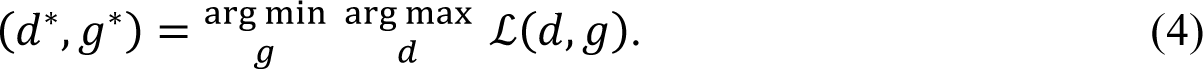

where 𝑑^∗^and 𝑔^∗^represent the trained critic and generator networks, respectively. Figure 1 provides an overview of the main blocks that make up the generator and critic, and Figure 2 shows these blocks in more detail.

**Figure 2.**
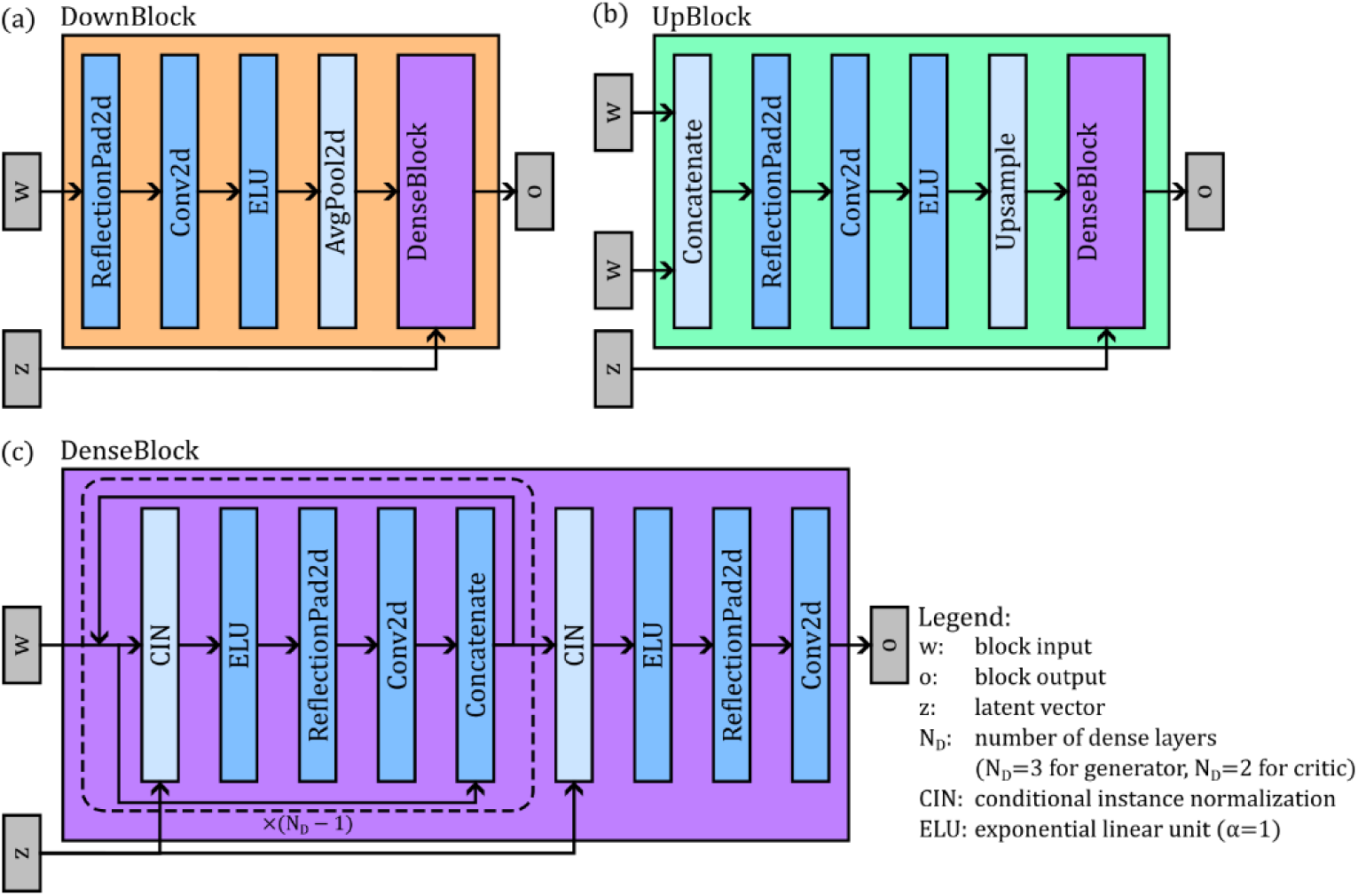
Building blocks that make up the cGAN. (a) Downblock, (b) UpBlock, and (c) DenseBlock.

### 2.3 Training and testing the cGAN

As described in Sec. 2.1.2, the training set for the cGAN consists of pairs of artificial elasticity images and phase difference images generated from FEA simulations. 5,000 artificial elasticity images were generated using a 2D parametric model representing a spheroid embedded in hydrogel. We used a 2D model and assumed plane strain as this greatly simplified the construction of the parametric model and reduced the computational complexity in the FEA simulation (thus allowing more training samples to be generated per unit time). Furthermore, since the experimental samples are loaded uniaxially, and deformation is constrained in the out-of-plane direction by adjacent material, the strain is predominantly along the vertical axis of the image, thus making the plane strain assumption reasonable. The spheroid was modelled as an ellipse that was subdivided by straight lines into cells, with each cell containing a circle representing the nucleus (as shown in Figure 3a). The boundaries between cells was determined by computing the 2D Laguerre-Voronoi diagram, or power diagram [54,55], which, for our purposes, allowed us to partition the spheroid into cells, each containing a nucleus located approximately at the center. Artificial elasticity images (Figure 3b) of dimensions 256 ×256 μm^2^ (256 ×256 pixels) were generated by randomly sampling the model parameters, namely, the spheroid size, spheroid height to width ratio, spheroid position, number of nuclei per unit area, nuclei positions, nuclei radii, nuclei elasticity, cytoplasm elasticity, and hydrogel elasticity. Poisson’s ratio for all components was set to 0.49 (i.e., all components were modelled as close to incompressible). The positions and sizes of nuclei were determined such that the minimum spacing between nuclei was 2 μm. The model parameters were sampled from a normal distribution. To constrain the parameter values to a finite range, values more than three standard deviations from the mean were resampled. The minima and maxima of the parameter distributions were determined by analysis of preliminary spheroid OCM, CFM, and mechano-microscopy images, and are summarized in Table 1.

**Figure 3.**
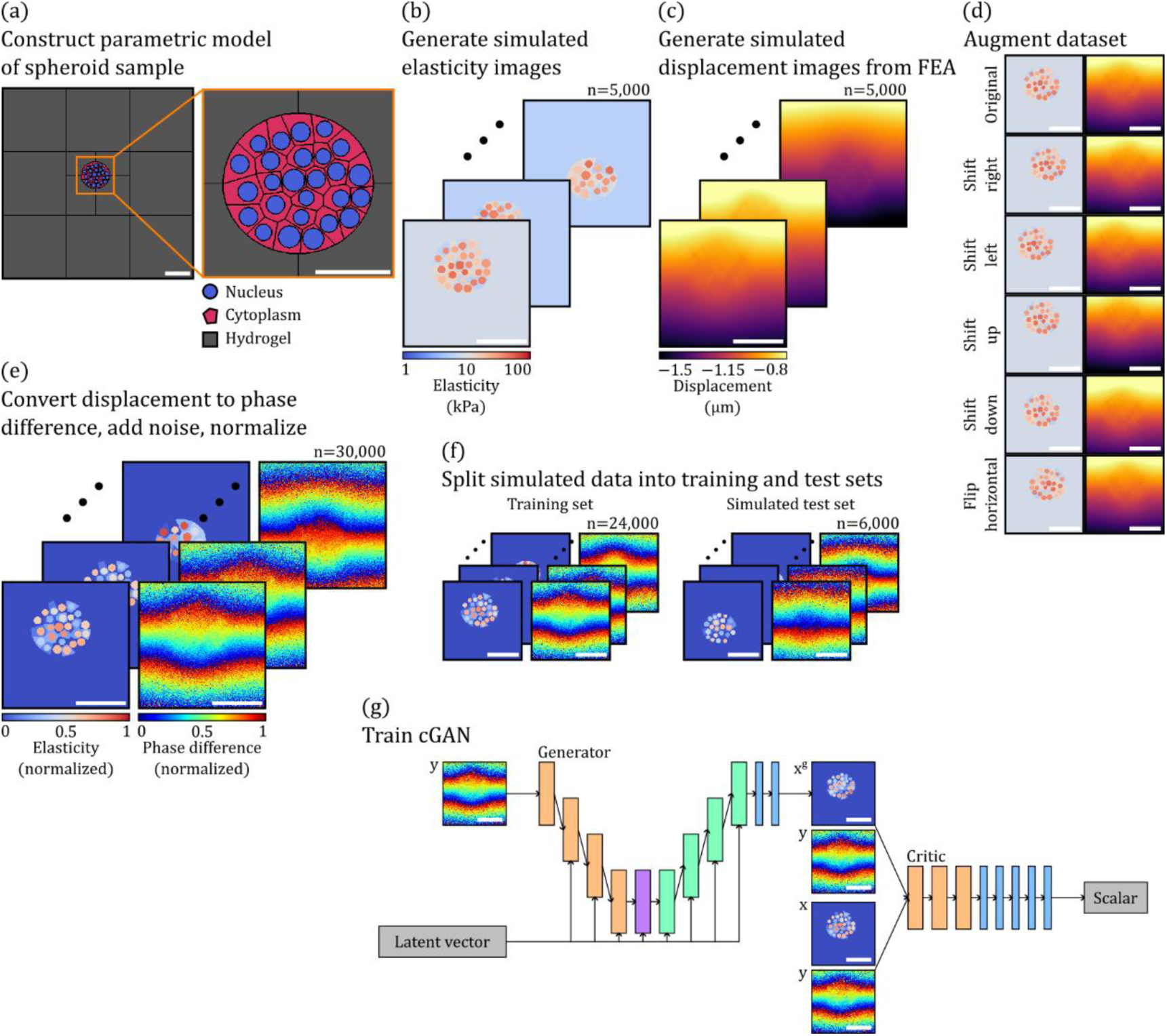
Overview of the steps required to train the cGAN, including: (a) constructing a parametric model of the spheroid samples; (b) generating the simulated elasticity images; (c) using FEA to compute the displacement images; (d) applying dataset augmentation to increase the dataset size; (e) preprocessing the data before input into the cGAN; (f) assembling the training set and simulated test set; and (g) training the cGAN. Scale bars: 100 μm.

**Table 1.** Distribution parameters from which the artificial elasticity image parameters were sampled.

| Parameter | Units | Minimum | Maximum |
| --- | --- | --- | --- |
| Spheroid width | $\mu\text{m}$ | 80 | 130 |
| Spheroid height to width ratio |  | 0.85 | 0.95 |
| Spheroid horizontal position | $\mu\text{m}$ | -40 | 40 |
| Spheroid vertical position | $\mu\text{m}$ | -40 | 40 |
| Number of nuclei per square millimeter |  | 4,000 | 5,500 |
| Nuclei radii | $\mu\text{m}$ | 6 | 9 |
| Nuclei Young's modulus | kPa | 15 | 50 |
| Cytoplasm Young's modulus | kPa | 5 | 15 |
| Hydrogel Young's modulus | kPa | 5 | 10 |
| Displacement applied to top edge | $\mu\text{m}$ | -4 | -0.4 |

The phase difference images corresponding to the artificial elasticity images were generated by first generating displacement images (Figure 3c) by simulating uniaxial compression using FEA software. During simulation, the artificial spheroid samples were compressed uniaxially. This was implemented by keeping the bottom-left corner fixed (to prevent rigid-body motion) while displacing the top edge downwards. As such, samples were free to expand laterally with no friction (this was approximated experimentally using lubrication, as discussed in Sec. 2.5). The amount of displacement applied to the top edge was sampled from a normal distribution in a manner similar to that used to obtain the model parameters, described previously, with the minimum and maximum shown in Table 1. The range of applied displacements was such that the corresponding strain was between 0.5 and 5 millistrain, similar to the range of strains seen in experimental mechano-microscopy data. Due to the low levels of strain applied, all materials were modelled as linear elastic. To reduce edge effects, the 256 ×256 μm^2^ artificial elasticity images were padded to 800 ×800 μm^2^. The mesh size was set to 1 μm in the central region and gradually increased to 10 μm at the outside edges of the padded regions. Structured tri meshing was used in the padded region, structured quad meshing was used in the central hydrogel region, and unstructured quad meshing was used in the spheroid. After simulation, the local axial displacements corresponding to each pixel in the artificial elasticity image was interpolated from the results. Finally, dataset augmentation (Figure 3d) was performed to further increase the number of samples sixfold (resulting in 30,000 samples) by shifting 20 μm up, down, left, and right, and flipping horizontally. All simulations were performed using Abaqus/Standard 2020 (Dassault Systèmes, Johnston, RI, USA) on a processing server running Windows Server 2012 R2 Standard with two 2.60 GHz Intel Xeon E5-2690 v4 octa-core central processing units (CPUs) and 192 GiB random access memory (RAM). No graphical processing unit (GPU) was available on this server. Each simulation took on average 92.9 seconds to complete.

The elasticity and displacement images required preprocessing prior to input to the cGAN (Figure 3e). Elasticity images were normalized using the formula 𝐸_𝑛_(𝑥, 𝑧) = log_10_(𝐸(𝑥, 𝑧)/𝐸_ℎ𝑦𝑑𝑟𝑜𝑔𝑒𝑙_), where 𝐸_𝑛_(𝑥, 𝑧) is the normalized elasticity at a given (𝑥, 𝑧) location, 𝐸(𝑥, 𝑧) is the unnormalized elasticity, and 𝐸_ℎ𝑦𝑑𝑟𝑜𝑔𝑒𝑙_ is the simulated hydrogel elasticity for a given sample, which scaled the images approximately to the range [0,1] while maintaining contrast between different components (i.e., nuclei, cytoplasm, hydrogel) in the images. As described in Sec. 2.1.2, displacement images were converted to phase difference images prior to input to the cGAN to avoid phase unwrapping artifacts. This was done by first converting the displacement images to unwrapped phase difference, Δ𝜙, using the equation 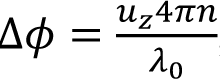, obtained by rearranging Eq. 1. To account for noise in the experimental phase difference images, noise was added to the simulated phase difference images used for training. In OCM images, the noise level is proportional to the SNR, and in a B-scan, the average SNR is not constant — instead, the SNR generally decreases with distance from the focus depth. For each depth, phase difference noise was assumed to be distributed according to the probability density function [56]

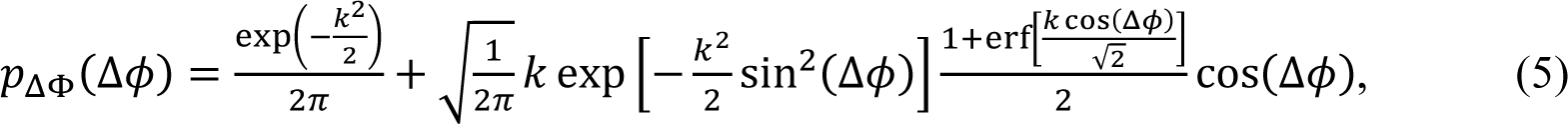

where 𝑘 is related to the phase difference variance 𝜎^2^, which is assumed to vary with depth along the z axis. The phase difference variance was determined as a function of 𝑧 (in millimeters) by calculating the variance at each depth in a region of homogeneous hydrogel and fitting the variance to the function

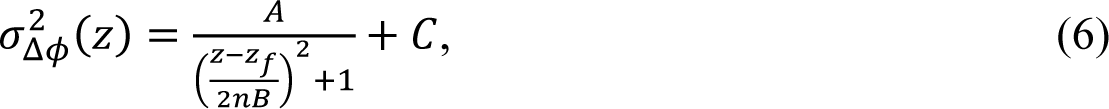

where 𝐴 = −2.764, 𝐵 = 0.06022, and 𝐶 = 3.170 are the fit parameters, 𝑧_𝑓_ is the focus depth, and 𝑛 = 1.4 is the refractive index of the hydrogel [11,32]. The focus depth was sampled from a uniform distribution with a minimum and maximum of 100 μm and 150 μm, respectively. Using Eqs. 5 and 6, a phase difference noise realization, 𝑁_Δ𝜙_, was generated and added to the phase difference, Δ𝜙_𝑁_ = Δ𝜙 + 𝑁_Δ𝜙_. The wrapped phase difference, Δ𝜙_𝑤_, was then computed using Δ𝜙_𝑤_ = ∠(exp[𝑖Δ𝜙_𝑁_]), such that −𝜋 < Δ𝜙_𝑤_ ≤ 𝜋 . Finally, the normalized phase difference, was computed using 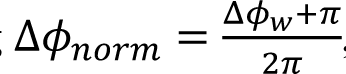, such that 0 < Δ𝜙

Of the 30,000 samples, 80% (24,000) were assigned to the training set, and 20% (6,000) were assigned to the test set (Figure 3f). All code for generating the datasets was written in Python 3.9.12, and the cGAN was implemented using PyTorch 2.0.1 [57]. The cGAN was trained for 200 epochs using a batch size of 16 and an Adam optimizer with a learning rate of 0.001, 𝛽_1_ = 0.5, 𝛽_2_ = 0.9, and weight decay parameter of 10^-7^ for both the generator and critic. Training took ∼76 hours using a high-performance computing cluster with an NVIDIA V100 Tensor Core GPU with 32 GB memory, a 2.10 GHz Intel Xeon Gold 6130 CPU, and 184 GB RAM.

The cGAN was evaluated using both simulated samples (referred to as the simulated test set) and real spheroids (referred to as the experimental test set). Since the cGAN generates a distribution of elasticity images for a given phase difference image (from which elasticity image instances may be sampled by varying the latent vector), the cGAN mean elasticity was computed using 30 instances for each phase difference image. A single cGAN elasticity image instance was produced in ∼1 second using the same machine used for training. For the simulated test set, the cGAN mean elasticity images were compared to the ground truth elasticity images, whereas for the experimental test set, the cGAN mean elasticity images were compared to the algebraic elasticity images, since the ground truth elasticity images were unavailable. As the cGAN outputs normalized elasticity images (𝐸_𝑛_(𝑥, 𝑧)), the outputs were unnormalized before comparison using the equation 𝐸(𝑥, 𝑧) = 𝐸_ℎ𝑦𝑑𝑟𝑜𝑔𝑒𝑙_ × 10^𝐸𝑛(𝑥,𝑧)^ . For the simulated test set, 𝐸_ℎ𝑦𝑑𝑟𝑜𝑔𝑒𝑙_ is known, whereas for the experimental test set, 𝐸_ℎ𝑦𝑑𝑟𝑜𝑔𝑒𝑙_ was estimated from the average elasticity computed using the algebraic method in a 50 × 50 μm^2^ (50 × 50 pixels) region in hydrogel, which was located using the OCM B-scan.

To evaluate the cGAN method and compare it to the algebraic method, the root mean squared error (RMSE) was computed for the simulated training and test sets using

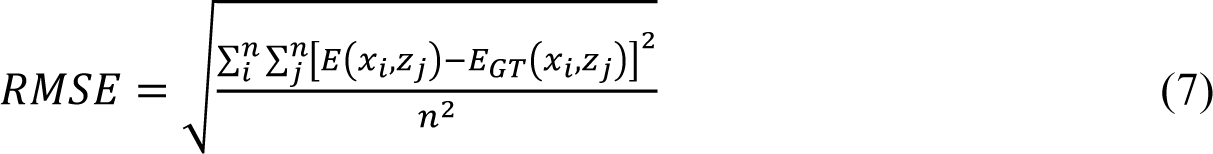

where 𝐸_𝐺𝑇_(𝑥_𝑖_, 𝑧_𝑗_) is the ground truth elasticity, 𝐸(𝑥_𝑖_, 𝑧_𝑗_) is the cGAN mean elasticity or the algebraic elasticity, and the summations are over all pixels in the images. As outlined in Sec. 2.1.1, the algebraic elasticity is usually computed by dividing the axial stress at the layer-sample interface by the axial strain in the sample. To apply the algebraic method to the simulated data, the axial stress at the layer-sample interface was approximated by extracting the axial stress from the bottom edge of the FEA simulation. The axial strain was computed using ordinary least squares regression over a 20 μm fitting range to calculate the gradient of the phase difference with respect to depth. To reduce the effect of noise, one-dimensional (1D) Gaussian filters with standard deviations of 2.4 μm and 0.3 μm were applied along the lateral dimension to the phase difference and strain images, respectively. Additionally, a median filter with a 3 × 3 pixel kernel was applied to the algebraic elasticity images to reduce the influence of small regions of very high elasticity that occur where the strain is very close to zero. For consistency, a median filter with the same kernel was also applied to the cGAN elasticity images.

### 2.4 Spheroid preparation

To determine suitable ranges for the parameter distributions used to construct the artificial spheroid samples, and to provide data to validate the trained cGAN, samples consisting of cell spheroids embedded in hydrogel were imaged using mechano-microscopy. To make the hydrogel, lyophilized gelatin methacryloyl (GelMA) was first placed in a desiccator to remove moisture, then dissolved in sterile Dulbecco’s Phosphate Buffered Saline (DPBS; Gibco), resulting in a 9% (w/v) GelMA solution. A photoinitiator, Irgacure-2959 (2-Hydroxy-4’-(2-hydroxyethoxy)-2-methylpropiophenone; Sigma-Aldrich), was then dissolved in absolute ethanol and added to the GelMA solution, resulting in a final concentration of 0.1% (w/v) Irgacure-2959. The solution was protected from light and stored at 4°C until cell encapsulation. MCF7 (non-metastatic, epithelial breast cancer) cells were cultured in an incubator using a media consisting of high glucose Dulbecco’s Modified Eagle Medium (hg-DMEM; Gibco), 1% (v/v) antibiotic-antimycotic (anti-anti; Gibco), and 10% (v/v) fetal bovine serum (FBS; Gibco). Before encapsulation, the GelMA solution was warmed to 37°C in a water bath for one hour. MCF7 cells were then added to the GelMA solution at a concentration of 2,000 cells/hydrogel. This solution was then pipetted into circular 5 mm diameter, 300 μm thick molds (polydimethylsiloxane; Wacker Chemie AG) placed on a glass slide treated with dichlorodimethylsilane (Sigma-Aldrich) and covered with a methacrylated (3-(trimethoxysilyl)propyl methacrylate; Sigma-Aldrich) circular 10 mm diameter coverslip. The samples were cured by exposure to UV light (365 nm, 2.5 mW/cm^2^) for 15–60 seconds and cultured for 7–16 days in a 12-well plate in normal growth medium (the same as that described previously) at 37°C 5% CO2 to enable spheroid growth. To enable CFM imaging of the live cells, the samples were stained within two hours of imaging. This was done by creating a dye mix by adding CellMask Green Plasma Membrane Stain (C37608; Thermo Fisher) to 0.5 mL of growth medium in 1:200 dilution and adding Hoechst 33342 Nucleic Acid Stain (62249; Thermo Fisher) in 10 μM final concentration. The growth medium in the 12-well plate was replaced with the dye mix (with the samples washed before the addition of the dye) and the samples were incubated at 37°C for 20 minutes. Finally, the samples were washed twice with warm PBS and returned to normal growth medium.

### 2.5 IMAGING

#### 2.5.2 **Mechano-microscopy**

To validate the outputs from the cGAN, samples were imaged using a mechano-microscopy system described previously [33] and summarized briefly here. The system is based on OCM, which is a high-resolution variant of OCT that uses high numerical aperture objectives to achieve high lateral resolution [32]. The light source consists of a supercontinuum laser (SuperK Extreme EXW-4 OCT; NKT Photonics) that is filtered to produce light with a spectral range of 650 nm to 950 nm (central wavelength of 800 nm), resulting in a measured OCM full width at half maximum (FWHM) axial resolution of ∼1.4 μm in air. In the sample arm, a two-axis galvanometer scanning system (GVSM002-EC/M; Thorlabs Inc) was used to scan the beam laterally, and a 20× 0.75 NA objective lens (CFI Plan Apo; Nikon) was used to focus the beam into the sample, resulting in a measured FWHM lateral resolution of ∼0.5 μm. A dual-arm configuration was used, with the reference arm containing the same optics as the sample arm to match optical dispersion, followed by a retroreflector (PS974M-B; Thorlabs Inc). The combined light from the sample and reference arms was detected by a spectrometer comprising a 2048-pixel line camera with a maximum line rate of 130 kHz (Wasatch Photonics).

To perform mechano-microscopy, after the objective lens, the sample arm contained a stack comprising, from top to bottom, a coverslip as an imaging window, a 300 μm thick GelMA sample with spheroids, a 1 mm thick compliant silicone layer, and a rigid glass fixed to an annular piezoelectric actuator (Piezomechanik GmbH). The sample and compliant layer were lightly compressed between the coverslip and rigid glass to ensure good contact between all components. OCM B-scans were acquired at 49 Hz and were synchronized with the actuator, which was driven by a 24.5 Hz square wave to apply loading to the sample, such that two B-scans were acquired at each 𝑦 location, one each for the unloaded and loaded states, respectively. Samples were scanned over a lateral field-of-view of 0.3 × 0.3 mm^2^, corresponding to 1,000 × 1,000 pixels.

#### 2.5.2 Confocal fluorescence microscopy

To visualize the subcellular morphology and function of the cell spheroids, a CFM system was integrated with the mechano-microscopy system. Using this setup, CFM images could be acquired of live cells following mechano-microscopy imaging without moving the sample, thus facilitating co-registration between CFM, OCM, and elasticity images. The CFM system consisted of an excitation light source (Changchun New Industries Optoelectronics Technology Co.) and photomultiplier tube (PMT2101/M; Thorlabs Inc.) as the detector. Two excitation wavelengths, 405 nm and 488 nm, were used to stimulate emission at 445 nm in the Hoechst 33342 Nucleic Acid Stain, and at 525 nm in the CellMask Green Plasma Membrane Stain, respectively. CFM and OCM were combined using a long-pass dichroic mirror (DMLP650; Thorlabs Inc.), such that both systems used the same sample arm galvanometer scanning system and optics, and the focal planes of both systems were aligned with each other. During CFM, the sample was also translated along the 𝑧 axis in steps of 5 μm to create a *z*-stack. The CFM axial and lateral resolutions were measured to be ∼8 μm and ∼0.5 μm, respectively.

## 3 Results

### 3.1 Simulated test set

Figure 4 shows box plots of the RMSE for the cGAN method applied to the training set, the cGAN method applied to the test set, and the algebraic method applied to the test set. To assess the relative contribution of the hydrogel region to the error versus that of the spheroid, the RMSE was computed for both the whole images (Figure 4a) and excluding the hydrogel regions (Figure 4b). To ensure independence between samples, sample augmentations (described in Sec. 2.3 and Figure 3d) were removed prior to plotting the box plots and performing statistical tests (the Wilcoxon signed-rank test) to assess the significance of the difference between the medians. From Figure 4a, the whole image RMSE for the cGAN method applied to the simulated test set (median: 3.47 kPa, 95% confidence interval (CI) [3.41, 3.52]) is not significantly different to that for the cGAN method applied to the training set (median: 3.50 kPa, 95% CI [3.48, 3.53]), indicating the network generalizes well and is not overfitting. Further, the whole image RMSE for the cGAN method applied to the simulated test set is significantly lower than that for the algebraic method applied to the simulated test set (median: 4.91 kPa, 95% CI [4.85, 4.97]), indicating that for simulated data, the cGAN method produces more accurate elasticity images than the algebraic method. This trend also holds for the spheroid RMSE (Figure 4b): here, the median spheroid RMSE for the cGAN method applied to the training set, the cGAN method applied to the test set, and the algebraic method applied to the test set are 10.1 kPa, 95% CI [10.1, 10.2]; 10.1 kPa, 95% CI [10.0, 10.2]; and 13.4 kPa, 95% CI [13.4, 13.5], respectively. While the spheroid RMSE is higher than the whole image RMSE across all groups (since both methods have a lower RMSE in hydrogel than in the spheroid), this result indicates that the cGAN improves elasticity estimation not only in the hydrogel, but also in the spheroid.

**Figure 4.**
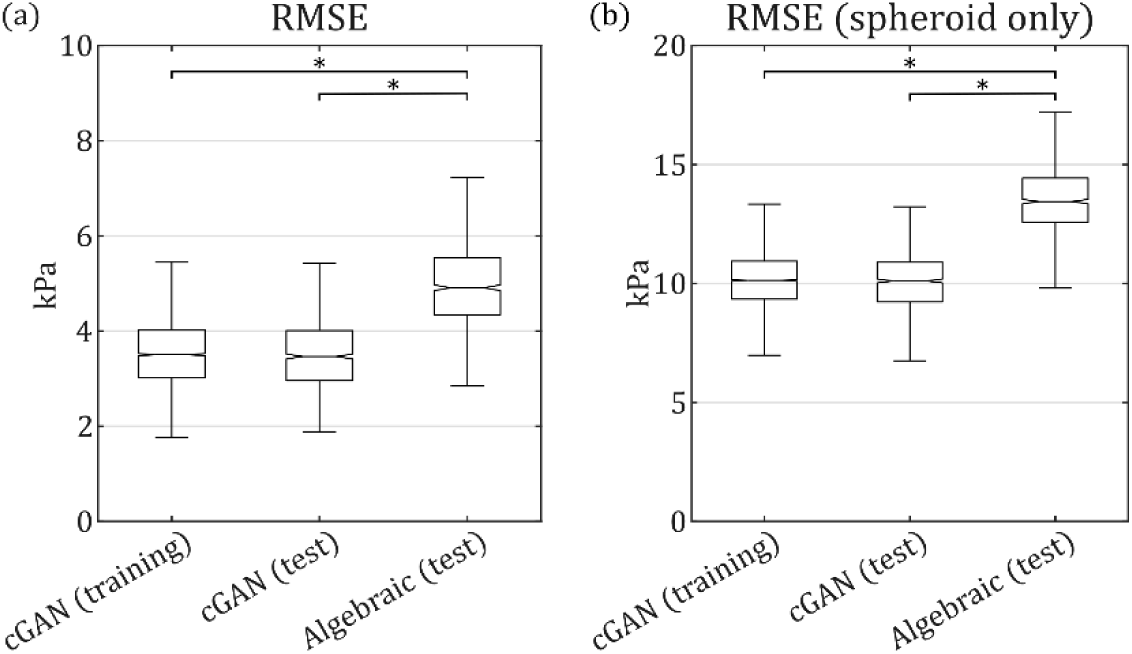
Box plots of RMSE between the simulated ground truth elasticities and the elasticities computed using the cGAN and the algebraic methods. For these plots, the cGAN method was applied to the unaugmented training set (n = 4000) and the unaugmented simulated test set (n = 1000), and the algebraic method was applied to the unaugmented simulated test set. This was done using (a) the entire elasticity images, and (b) only the spheroid regions of the elasticity images. For each box plot, the central mark indicates the median; the notch indicates the 95% confidence interval of the median approximated by ±1.57 × 𝐼𝑄𝑅/√𝑛, where IQR is the interquartile range; the bottom and top edges of the box indicate the 25^th^ and 75^th^ percentiles, respectively; and the bottom and top whiskers extend to the most extreme data points less than 1.5 times the interquartile range from the bottom and top edges of the box, respectively. Asterisks indicate a significant difference (p < 0.05) between the medians.

To illustrate why the mean squared error for the cGAN elasticities is lower than that for the algebraic elasticities, Figure 5 shows a representative sample from the simulated test set with the corresponding cGAN elasticity and algebraic elasticity. Comparing the ground truth elasticity (Figure 5d) with the cGAN mean elasticity (Figure 5b) and the algebraic elasticity (Figure 5e), we can see that the spheroid is visible against the hydrogel background in all three elasticity images. While the cGAN elasticity image does not perfectly replicate the ground truth elasticity image, there is nonetheless a close correspondence, with many of the nuclei visible in the ground truth image matched to a nucleus at the corresponding location in the cGAN elasticity image. Additionally, the cGAN elasticity image exhibits better resolution and contrast than the algebraic elasticity image. For example, the hydrogel elasticity appears more uniform in the cGAN elasticity image compared to the algebraic elasticity image, in agreement with the ground truth. Additionally, while some of the stiffer nuclei are faintly visible in the algebraic elasticity image (indicated by the yellow arrowheads), the softer nuclei are harder to locate (indicated by the teal arrowheads), however, in the cGAN elasticity image, all these nuclei are easily identified. The improved accuracy and resolution of the cGAN elasticity can also be seen from the line plot of elasticity and strain versus depth (Figure 5i), which shows that the cGAN elasticity matches the true elasticity better than the algebraic elasticity. An additional benefit of the cGAN method is that uncertainty quantification of the cGAN elasticity is provided by the cGAN elasticity standard deviation (Figure 5c), where lower standard deviation indicates lower uncertainty. Figure 5c shows that the lowest uncertainty is in the hydrogel, while high uncertainty tends to be present at the edges of nuclei, likely due to the sharp change in elasticity at these regions. Within the spheroid, the cGAN tends to be least uncertain at the center of nuclei and in cytoplasm; in particular, the two regions indicated by the magenta arrowheads correspond to cytoplasm regions with especially low uncertainty in elasticity, for which the cGAN elasticity appears to closely match the ground truth elasticity. The poor visibility of the nuclei in the algebraic elasticity image can be explained by considering that the algebraic elasticity image is obtained by dividing the axial stress extracted from the bottom edge of the FEA simulation (Figure 5h) by the axial strain (Figure 5f). As discussed in Sec. 2.1.1, the algebraic method assumes the stress is uniaxial and axially uniform (Figure 5h), however, by viewing the axial stress throughout the sample computed by FEA (Figure 5g), we see this is not the case, as mechanical heterogeneity in the sample causes regions of stress concentration. Therefore, the algebraic method generally underestimates the magnitude of the stress within the nuclei, resulting in underestimation of the nuclei elasticity. Additionally, the strain image (Figure 5f) shows that strain is highly sensitive to noise in the phase difference image (as demonstrated by the heterogeneity of the strain in the hydrogel); furthermore, the strain resolution is reduced by Gaussian filtering and the need to compute the strain by linear regression over a fitting range. We also note that the prior information embedded in the cGAN method, through the selection of the training samples, further helps it recover elastic modulus fields that are consistent with a typical spheroid. The combination of these factors results in poorer contrast and resolution in the algebraic elasticity image compared to the cGAN elasticity image.

**Figure 5.**
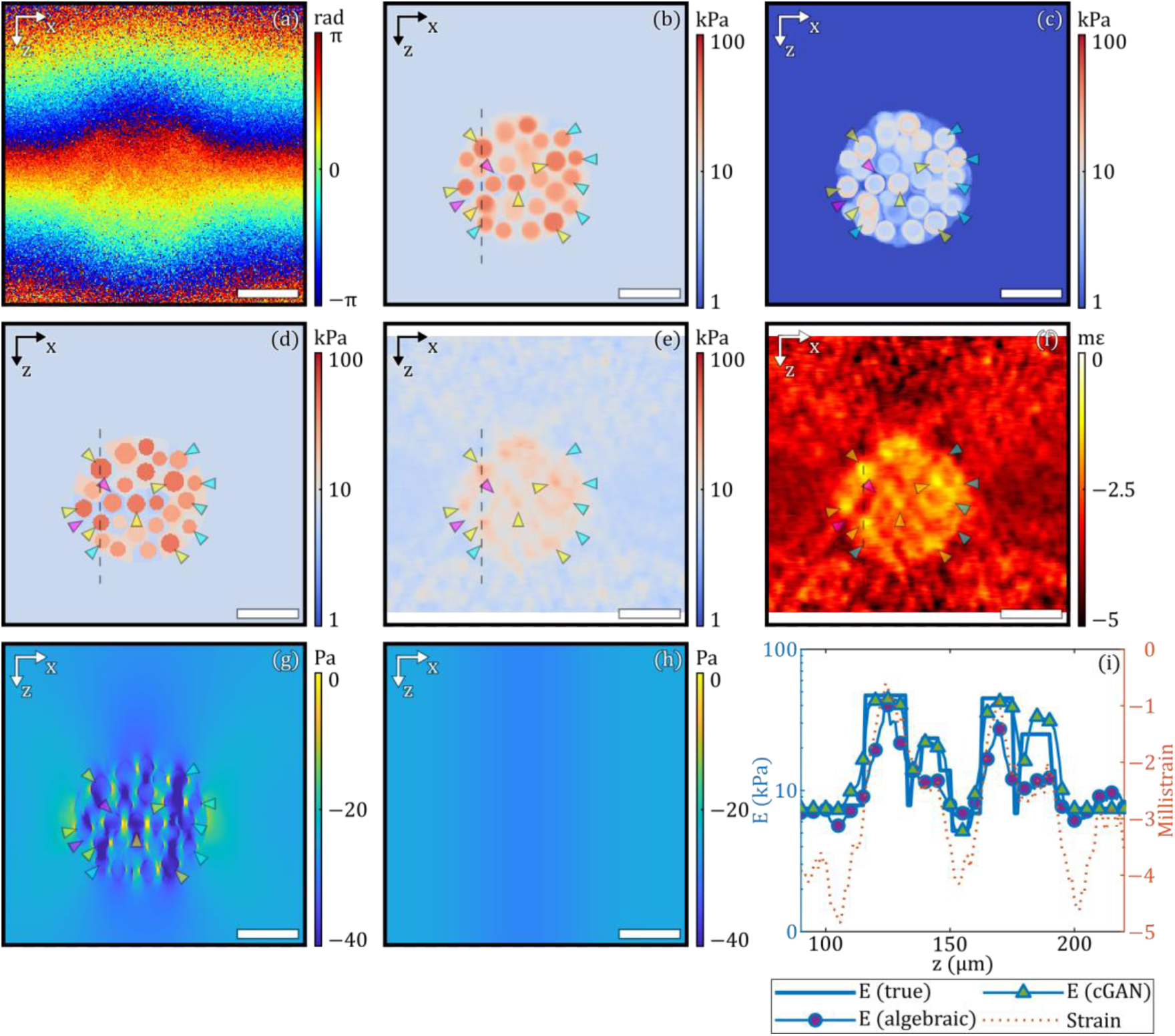
A representative sample from the simulated test set. (a) Simulated phase difference, (b) cGAN mean elasticity, (c) cGAN elasticity standard deviation, (d) simulated (ground truth) elasticity, (e) algebraic elasticity, (f) axial strain, (g) axial stress from FEA, (h) axial stress extracted from the bottom edge of (g) representing the layer stress in the algebraic method, and (i) line plots of elasticity and strain versus depth corresponding to the dashed lines in (b) and (d)–(f). Yellow arrowheads indicate circles with elevated elasticity in the cGAN elasticity image that correspond to elevated elasticity locations in the algebraic elasticity image. Blue arrowheads indicate circles with elevated elasticity in the cGAN elasticity image that are difficult to locate in the algebraic elasticity image. Magenta arrowheads indicate regions with low cGAN standard deviation corresponding to low uncertainty in the cGAN output. Scale bars: 50 μm.

### 3.2 Experimental test set

To assess the cGAN on real data, we applied it to phase difference measurements obtained experimentally by imaging samples containing tumor spheroids embedded in a hydrogel. Two examples are shown in Figure 6 and Figure 7. Although the ground truth elasticity is unknown for the experimental data, thus precluding direct evaluation of the cGAN elasticity images by comparison with the ground truth, we present co-registered algebraic elasticity images, as well as OCM and CFM images to help identify the biological structures that correspond to features in the elasticity images. Additionally, we present axial strain images to aid the explanation of features seen in the algebraic elasticity images. Note that while the phase difference, elasticity, OCM, and strain images are in the same plane (the B-scan plane), the CFM image is in an orthogonal plane (the *en face* plane). This is because in Abaqus we must simulate the B-scan plane, since the boundary conditions (i.e., displacements) are known at the top and bottom of the B-scans, which results in the cGAN also generating elasticity images in the B-scan plane. However, the CFM images are displayed in the *en face* plane because the CFM system uses the same sample arm optics as the OCM system, such that the CFM images exhibit much better resolution in the *en face* plane compared slicing the *z*-stack in the B-scan plane.

**Figure 6.**
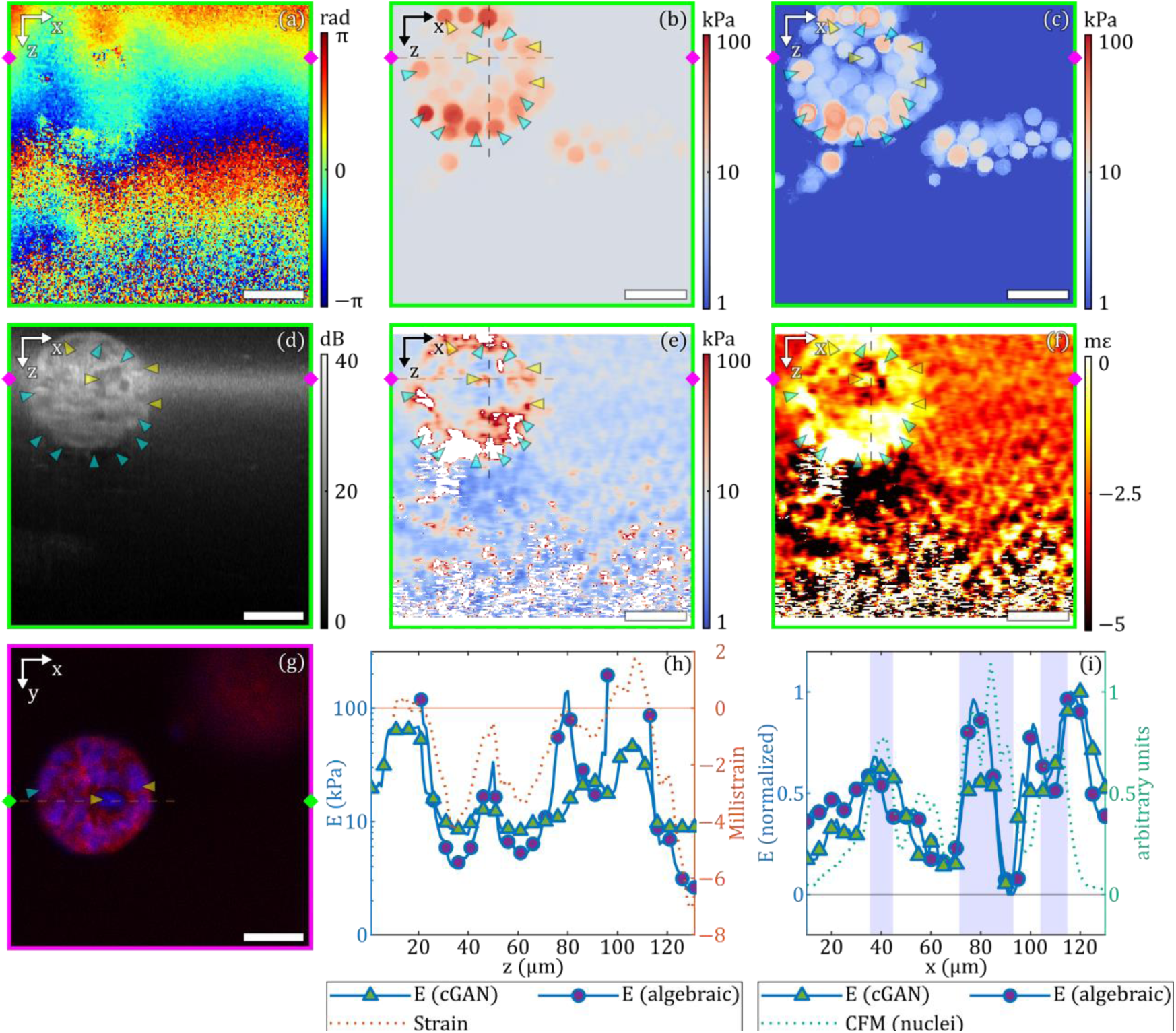
First example from the experimental test set. (a) Experimental phase difference, (b) cGAN mean elasticity, (c) cGAN elasticity standard deviation, (d) OCM, (e) algebraic elasticity, (f) axial strain, (g) CFM, (h) line plots of elasticity and strain versus depth corresponding to the black dashed lines in (b), (e), and (f), and (i) line plots of elasticity and nucleic acid stain intensity versus x corresponding to the brown dashed lines in (b), (e), and (g). Subfigures (a)–(f) show the B-scan plane at the y location indicated in (g) by the green diamonds, and subfigure (g) shows the en face plane at the z location indicated in (a)–(f) by the magenta diamonds. Yellow arrowheads indicate circles with elevated elasticity in the cGAN elasticity image that correspond to elevated elasticity locations in the algebraic elasticity image, dark regions in the OCM image and nuclei in the CFM image. Blue arrowheads indicate circles with elevated elasticity in the cGAN elasticity image that correspond to negative elasticity in algebraic elasticity image and dark regions in the OCM image. Blue rectangles in (i) indicate locations where the nucleic acid stain intensity is greater than 0.5. Scale bars: 50 μm.

**Figure 7.**
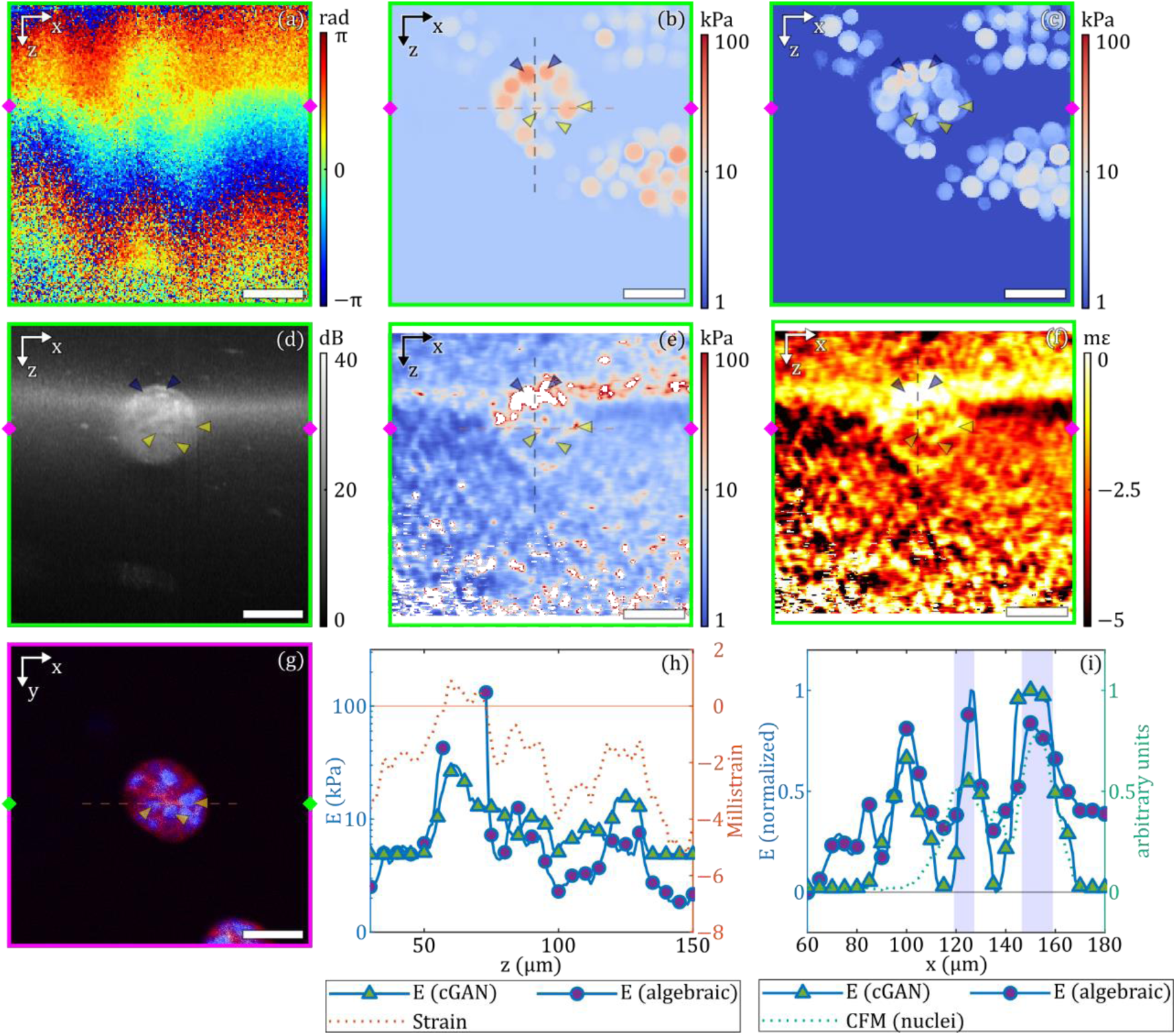
Second example from the experimental test set. (a) Experimental phase difference, (b) cGAN mean elasticity, (c) cGAN elasticity standard deviation, (d) OCM, (e) algebraic elasticity, (f) axial strain, (g) CFM, (h) line plots of elasticity and strain versus depth corresponding to the black dashed lines in (b), (e), and (f), and (i) line plots of elasticity and nucleic acid stain intensity versus x corresponding to the brown dashed lines in (b), (e), and (g). Subfigures (a)–(f) show the B-scan plane at the y location indicated in (g) by the green diamonds, and subfigure (g) shows the en face plane at the z location indicated in (a)–(f) by the magenta diamonds. Yellow arrowheads indicate circles with elevated elasticity in the cGAN elasticity image that correspond to elevated elasticity locations in the algebraic elasticity image, dark regions in the OCM image and nuclei in the CFM image. Blue arrowheads indicate circles with elevated elasticity in the cGAN elasticity image that correspond to negative elasticity in algebraic elasticity image and dark regions in the OCM image. Blue rectangles in (i) indicate locations where the nucleic acid stain intensity is greater than 0.5. Scale bars: 50 μm.

Figure 6 shows a sample containing a spheroid in the top-left corner. Overall, the cGAN mean elasticity image (Figure 6b) and the algebraic elasticity image (Figure 6e) appear consistent, with both showing regions of high elasticity corresponding to the spheroid location shown in the OCM image (Figure 6d). In both elasticity images, the spheroid appears to contain more stiff nuclei at the periphery than at the center, which is consistent with previous literature [11]. In the hydrogel region, the cGAN elasticity image generally appears less noisy than the algebraic elasticity image (especially towards the bottom of the images), however, a cluster of high-elasticity circles are visible in the cGAN elasticity image that are outside the spheroid (near the bottom-right side of the spheroid), which may have been caused by noise. Additionally, several nuclei in the cGAN elasticity image correspond with elasticity peaks in the algebraic elasticity image, as well as dark regions in the OCM image (indicated by yellow arrowheads), and peaks in the nucleic acid stain channel of the CFM image (Figure 6i) which suggests that these nuclei in the cGAN elasticity image are indeed real. Note that in Figure 6i (and Figure 7i), the cGAN and algebraic elasticities have been normalized by taking the base ten logarithm and applying linear scaling such that the minimum and maximum of each plot to zero and one, respectively. This is done to highlight the correspondence between peaks in the nucleic acid stain intensity, which is also shown on a normalized scale, with peaks in the elasticity plots, which can be difficult to identify if displayed without normalization in cases where the cGAN and algebraic method peak elasticities differ greatly in magnitude. In Figure 6i, three peaks are visible in the nucleic acid stain intensity (indicated by the blue rectangles) that appear to correspond to three peaks in both the algebraic and cGAN elasticity plots. Interestingly, several nuclei in the cGAN elasticity image also seem to correspond to regions of negative elasticity in the algebraic image (indicated by blue arrowheads), which have been masked. These regions occur due to positive strain (see Figure 6f and Figure 6h), which may occur due to shear stresses, local compressibility within the spheroid, or noise in regions exhibiting low strain, which could be caused by stiff nuclei. Therefore, an advantage of the cGAN method may be to visualize stiff nuclei that induce positive strain, and therefore produce negative elasticity when using the algebraic method.

Figure 7 shows an example containing a spheroid slightly above the center of the image. Similar to the previous example, both the cGAN elasticity image (Figure 7b) and the algebraic elasticity image (Figure 7e) appear largely consistent, with both showing a circular region of high elasticity corresponding to the spheroid location (as seen in the OCM image in Figure 7d) that exhibits higher elasticity at the periphery. In this example the algebraic elasticity image features a horizontal line of high elasticity that intersects with the top of the spheroid, which is likely an artifact from linear fitting to obtain strain across the OCM focal plane (visible as the bright horizontal band in Figure 7d). This artifact is absent from the cGAN elasticity image, suggesting that the cGAN may be more robust to such artifacts, although, again, in the cGAN elasticity image, there are a few clusters of high elasticity circles outside the spheroid towards the top-left, top-right, and bottom-right of the spheroid, which are likely the result of noise. Similar to the previous example, several nuclei in the cGAN elasticity image correspond to either elasticity peaks (yellow arrowheads) or negative elasticity (blue arrowheads) in the algebraic elasticity image; furthermore, these features correspond to dark regions in the OCM image (Figure 7d) and nuclei in the CFM image (Figure 7g), suggesting they correspond to real nuclei. In Figure 7i, three peaks are visible in the cGAN elasticity, which correspond to three peaks in the algebraic elasticity. While the leftmost peak (at x ≈ 100 μm) does not correspond to a peak in the nucleic acid stain intensity plot, the remaining two do, indicating these correspond to real nuclei.

## 4 Discussion

In this study, we developed a methodology to train a cGAN to generate elasticity images of samples containing a spheroid embedded in hydrogel from displacement measurements in the form of phase difference images. This involved building training and test sets from simulated data consisting of pairs of elasticity images and phase difference images, where elasticity images were generated from artificial samples built using a parametric model, and phase difference images were derived from displacement maps computed using FEA to simulate compression applied to the artificial samples. The cGAN was evaluated using simulated data by comparison with the known ground-truth elasticity, and experimental data from mechano-microscopy by comparison with elasticity images generated using the algebraic method, OCM images, and CFM images. For the simulated test set, our results indicate that the cGAN method produces elasticity images that represent the ground truth elasticity more accurately and with better resolution than the algebraic method. The improved accuracy is demonstrated by the reduced median RMSE of the mean cGAN elasticity images compared to the algebraic elasticity images (3.47 kPa, 95% CI [3.41, 3.52] versus 4.91 kPa, 95% CI [4.85, 4.97], respectively, for the whole images, and 10.1 kPa, 95% CI [10.0, 10.2] versus 13.4 kPa, 95% CI [13.4, 13.5], respectively, for the spheroid regions; see Figure 4). Visual comparison between the ground truth elasticity image and the mean cGAN elasticity image for an example test sample (Figure 5) also shows that the cGAN method can generate elasticity images that closely resemble the ground truth elasticity images; for example, the position, size, and elasticity of many of the nuclei generated in the mean cGAN elasticity image closely correspond to nuclei in the ground truth image. In contrast, while some of the larger and stiffer nuclei are faintly visible in the algebraic elasticity image, overall, the nuclei appear much sharper in the cGAN image. This is mainly because the algebraic method requires least squares regression over a fitting range (in this case, 20 μm) to compute axial strain. This process reduces axial resolution and is highly sensitive to noise, which necessitates spatial averaging that further reduces resolution. As this process is not required for the cGAN method, it is possible to generate elasticity images of spheroids with higher resolution using this method.

For the experimental test set, no ground truth elasticity is available to directly determine the accuracy of the cGAN elasticity images. Therefore, we compared the cGAN elasticity images with the algebraic elasticity images. In Sec. 3.1, we showed that stiff regions often appear as high elasticity peaks in algebraic elasticity images despite their limited resolution, and in Sec. 3.2 we identified several regions where the nuclei in the cGAN image coincide with high elasticity peaks in the algebraic elasticity image, suggesting that the cGAN image is correct in placing stiff nuclei in these regions. Additionally, we highlighted stiff regions in the cGAN elasticity image and corresponding features in the OCM and CFM images to provide further evidence for the presence of structures in these locations. We also identified regions where the algebraic method produced negative elasticity, likely due to noise in low-strain (i.e., stiff) regions or shear stresses pushing the sample in the opposite direction to compression. Many of these regions exhibit round shapes reminiscent of nuclei, which the cGAN method has likely correctly interpreted as stiff nuclei, thus demonstrating an advantage of the cGAN method over the algebraic method.

Ideally, to perform experimental validation of the cGAN method, we would apply the cGAN method to a real sample resembling a spheroid embedded in hydrogel with known elasticity (i.e., a spheroid-mimicking phantom). In theory, such a phantom could be fabricated by embedding silicone spheres, representing nuclei, within larger spheres, representing cells, in hydrogel, however, this is very difficult in practice due to the small length scales involved (e.g., ∼100 μm for the spheroid diameter). As the production of spheroid phantoms would require the development of new phantom manufacturing techniques, in this study, we performed experimental validation of the cGAN method by comparison with algebraic elasticity, OCM, and CFM images. Spheroid-mimicking phantom production may be an avenue for future research.

In this study, it was necessary to balance the accuracy of the FEA simulations with their complexity. For example, in this study, a 2D FEA model was used under the assumption of plane strain. While it may have been more realistic to take a 2D slice from a 3D FEA model, there is a tradeoff between the training set size and the model complexity, since a more complex model would take longer to simulate, resulting in fewer training samples per unit time. Additionally, the samples were assumed to consist only of nuclei, cytoplasm, and hydrogel, with each of these components being homogeneous and mechanically bonded to each other, however, cells contain structures within the nuclei and cytoplasm that were not considered in our model. Again, this simplification reduces the computational time required for FEA simulation (thus increasing the size of the training set that can be constructed) at the expense of accuracy. Future work could investigate the optimal balance between these tradeoffs.

Several possibilities exist as to the next steps for this work. Firstly, the methodology presented here (i.e., constructing a training set using FEA and using this to train a machine learning model to generate elasticity images conditional upon real data) is general enough that there may be a wide range of alternative suitable applications, once the application is simple enough to represent using a parametric model that enables the random generation of new samples. While we have demonstrated this methodology with a cGAN, alternative models may be substituted, such as conditional diffusion models, for which promising results have been demonstrated in computer vision, natural language generation, and medical image reconstruction [58]. In addition, physics-based models [59] may be employed to incorporate the laws of physics into the network training process, which may improve the quality of the outputs.

## 5 Conclusion

In this study, we trained a cGAN to generate elasticity images from phase difference images of samples containing a tumor spheroid embedded in a hydrogel and compared the outputs to elasticity images generated using the current signal processing chain in mechano-microscopy, the algebraic method. We evaluated the cGAN on both simulated data and experimental phase difference images of real spheroids and compared the cGAN elasticity with the algebraic elasticity, OCM, and CFM. Compared to the algebraic elasticity, the cGAN elasticity exhibits better spatial resolution, sensitivity, and robustness to noise, especially within stiff nuclei.

## Acknowledgements

This work was supported by the Cancer Council Western Australia (MSH, YSC, and BFK); the Australian Research Council DP220100163 (YSC and BFK); the Ian Potter Foundation (BFK); the Australian Government Research Training Program Scholarship (DV and SEA); the Hackett Postgraduate Research Scholarship (SEA); and ARO, USA grant W911NF2010050 (AAO). BFK acknowledges funding from the NAWA Chair programme of the Polish National Agency for Academic Exchange and from the National Science Centre, Poland. The authors gratefully acknowledge Dr. Liisa M. Hirvonen from the Centre for Microscopy, Characterisation & Analysis, The University of Western Australia, for assistance with live cell staining; and the Center for Advanced Research Computing (CARC) at the University of Southern California, USA, for providing computing resources that have contributed to the research results reported within this publication.

